# Combinatorial action of regulatory systems generates colistin heteroresistance

**DOI:** 10.1101/2025.08.11.668767

**Authors:** Jacob E. Choby, Emily K. Crispell, Muqing Ma, Linda M. Vu, Mary-Kate Key, Robert K. Ernst, Minsu Kim, David S. Weiss

## Abstract

Heteroresistance is a form of antibiotic resistance in which a minor subpopulation of resistant cells coexists with a majority susceptible population. Colistin heteroresistance is common among *Enterobacter cloacae* clinical isolates, threatens its utility as a last-line therapeutic, and has become a model with which to understand the fundamental bases of heteroresistance. Despite numerous insights, the mechanism by which phenotypic heterogeneity is generated within the population and leads to colistin heteroresistance has been unclear. Here, using a transposon-based mutagenesis screen, we identify the sigma factor σ^E^ as the source of heterogeneity in population-wide colistin resistance levels. Single-cell tracking experiments revealed that σ^E^ is active in only one percent of the population at baseline, and only those cells with active σ^E^ survive colistin exposure. However, σ^E^ expression and population heterogeneity are insufficient for survival, as a mutant lacking the PhoPQ two-component system controlling lipid A modifications necessary for colistin resistance retains heterogeneity but loses colistin resistance. These findings lead to a new paradigm in heteroresistance, where the combinatorial action of multiple regulatory systems, encompassing a heterogeneity generator and a distinct resistance generator, are required to give rise to colistin heteroresistance.

## INTRODUCTION

Colistin is a last-resort antibiotic reserved for treatment of multi-drug resistant Gram-negative bacteria, but its usefulness is challenged by emerging resistance. A member of the polymyxin class, colistin is a cationic antimicrobial peptide (CAMP) that targets the Gram-negative outer membrane constituent lipopolysaccharide (LPS) and disrupts the outer and inner membranes, killing the bacterium (*1*). Gram-negative bacteria can resist colistin killing by changing the composition of the outer membrane, in particular the lipid A portion of LPS (*2*). Generally, carbapenem-resistant Enterobacterales, a WHO Critical Priority (*3*) and CDC Urgent Threat (*4*), are designated colistin susceptible. However, conventional colistin resistance can occur when an isolate has intrinsic or horizontally-acquired LPS-modifying enzymes. Equally or more common, but challenging to detect (*5–8*), is a form of phenotypic colistin resistance termed heteroresistance (*9*).

Heteroresistance (HR) is a type of phenotypic heterogeneity in which cells in a population exhibit differences in relative antibiotic resistance; a low frequency subpopulation of cells is markedly more resistant than the majority susceptible population. The resistant subpopulation can be extremely rare (as few as 1 per 10^6^ cells) and is thus often undetected by conventional antimicrobial susceptibility testing. The subpopulation is also dynamic: the resistant cells are enriched in the presence of a given antibiotic but return to their baseline frequency in its absence. For these reasons, HR can contribute to failure of antibiotic therapy, when a patient is treated with an antibiotic to which the infecting organism exhibits undetected HR (*10–13*).

Colistin HR among carbapenem-resistant Enterobacterales is especially common in the *Enterobacter* genus (*5, 8*). The *Enterobacter* are diverse, and many pathogenic species are often collectively referred to as the *Enterobacter cloacae* complex (*14*). They are leading nosocomial, opportunistic pathogens and cause pneumonia, bloodstream infections, and urinary tract infections (*15*). A challenge for treating infections caused by *E. cloacae* complex isolates is the high rate of antibiotic resistance, especially to cephalosporins and carbapenems, making colistin an important last-line treatment option.

We previously described colistin HR in a cephalosporin-resistant *Enterobacter* clinical isolate (*16*). That work, and analyses of other colistin HR *Enterobacter* isolates, have resulted in colistin HR serving as a model for Gram-negative antibiotic HR. The resistant subpopulation in colistin HR *Enterobacter* harbors 4-amino-4-deoxy--arabinose (ʟ-Ara4N) modifications on the lipid A moiety of LPS, increasing cellular surface charge and thus reducing binding of cationic colistin and other CAMPs (*8, 16–19*). The *arnBCADTEF* gene cassette encodes the ʟ-Ara4N modification machinery and its expression is activated by the two-component system PhoPQ (*2*). Different alleles of *phoP*, *phoQ,* their promoter (*8, 17*), the PhoPQ-regulating *mgrB* (*8*), or *ecr* (*20*), affect PhoPQ/*arn* activation and the subsequent frequency of the resistant subpopulation. However, the underlying basis for the phenotypic heterogeneity exhibited within the cellular population of a colistin HR isolate has remained unresolved.

In this work, we used a transposon-based mutagenesis screen and single-cell tracking to identify the alternative sigma factor σ^E^ as the critical determinant of phenotypic heterogeneity in colistin HR in an *Enterobacter* clinical isolate. σ^E^ responds to envelope stress and activates transcription of a large stress response regulon (*21*). We observed that the resistant subpopulation had distinctly high σ^E^ activity which resulted in those cells surviving colistin treatment. σ^E^ increased ʟ-Ara4N modification of lipid A and responded to antimicrobial peptide exposure. However, while σ^E^ was the critical determinant of phenotypic heterogeneity, it was insufficient to cause HR, since a *phoPQ* mutant strain maintained heterogeneity but lost HR. Thus, we have identified σ^E^ as the generator of heterogeneity in the population that, in tandem with the resistance enhancement conferred by PhoPQ and the Arn machinery, establishes a rare, colistin resistant subpopulation. This establishes a novel mechanistic paradigm where the distinct but combinatorial activity of two regulatory systems generates heteroresistance, an enigmatic yet increasingly recognized source of antibiotic failure in the clinic.

## RESULTS

### Colistin heteroresistance requires lipid A modification

*Enterobacter cloacae* strain RS, a bloodstream isolate from a renal transplant patient, exhibits colistin HR and caused colistin treatment failure in a murine model of infection (*16*). Population analysis profile (PAP; Supp. Figure 1), the gold standard test to sensitively detect HR, which involves plating a range of dilutions of a bacterial strain at a range of concentrations of a given antibiotic, revealed that ∼1% of the total RS population (−2 log) was resistant to 4 μg/mL colistin, the CLSI breakpoint concentration used to discriminate susceptible from resistant strains. In contrast, the ColR2 strain demonstrated homogeneous resistance; every cell in the population survived at high colistin concentrations. ColS1 was susceptible to the colistin breakpoint concentration with no cells surviving (Figure 1a, Supp. Figure 2). When exposed to colistin, the resistant subpopulation of cells in strain RS survived and were enriched (Figure 1b). However, characteristic of HR, when colistin was removed, the frequency of the resistant subpopulation returned to the baseline level within one passage, demonstrating that the phenotypically resistant subpopulation can be transiently enriched and is not the result of stable genetic changes that confer resistance (Figure 1b). Additionally, when the genome of the resistant subpopulation was compared to that of the majority susceptible population, no sequence variants were detected (*16*) and no gene copy number variation was observed (Supplemental Table 1), consistent with phenotypic differences between the resistant and susceptible cells that are not due to genetic changes.

**Figure 1.**
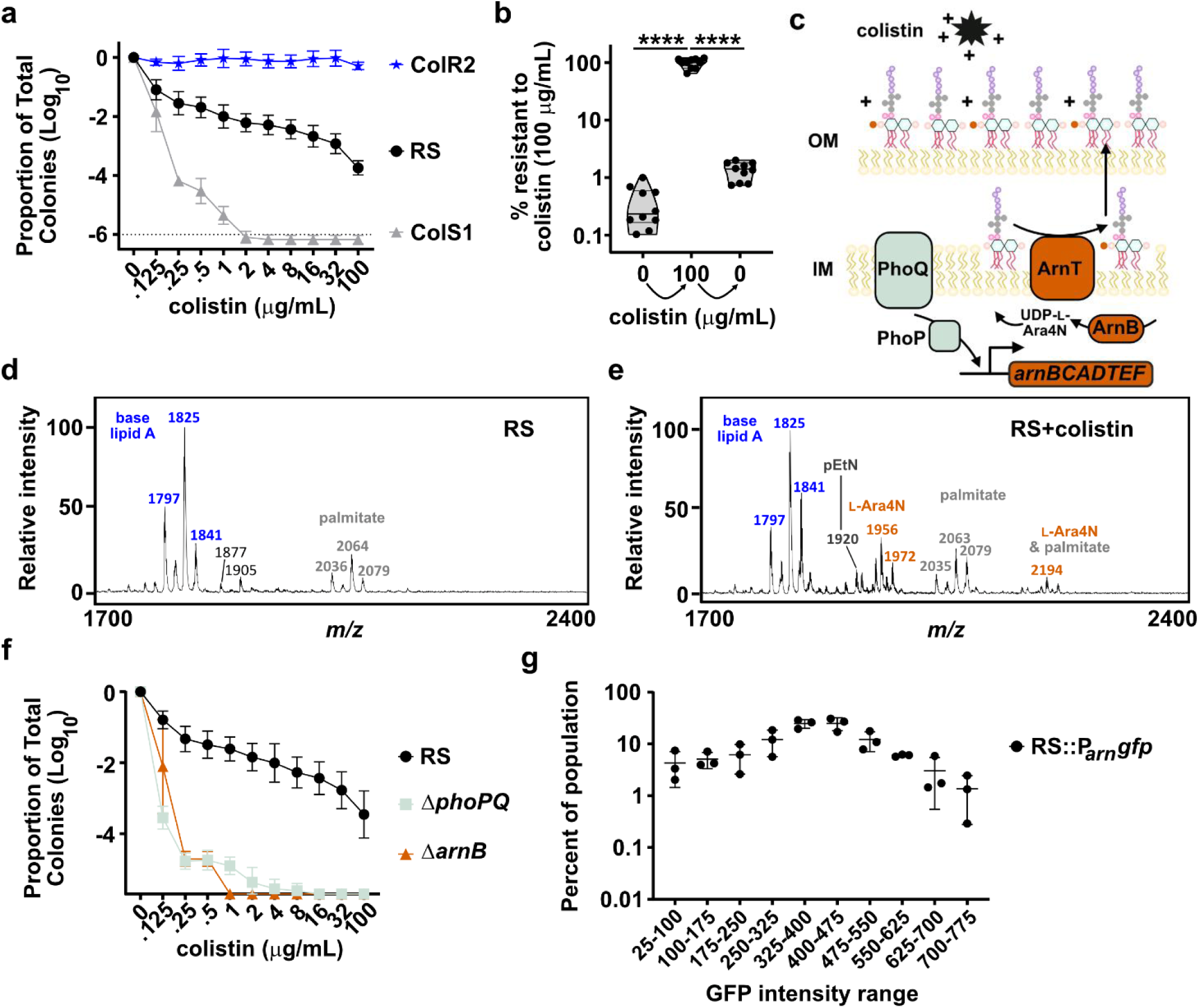
4-amino-4-deoxy-ʟ-arabinose modification of lipid A is required for the resistant subpopulation in *Enterobacter cloacae* RS. (a) Population analysis profile (PAP) of strains RS, ColS1, and ColR2 plated on Mueller-Hinton agar (MHA) containing colistin; the proportion of surviving colonies is quantified relative to MHA containing no colistin, from three independent experiments with n=8 total biological replicates. (b) Quantification of the colistin resistant subpopulation of RS in media alone to exponential phase (0), subcultured into 100 μg/mL colistin and grown for 22 h (100), and subcultured into fresh media without colistin and grown to exponential phase (0). At each point, an aliquot was diluted and plated onto MHA containing 100 μg/mL colistin to quantify the resistant subpopulation, from two independent experiments with n=10 total biological replicates. **** indicates p<0.0001 by RM one-way ANOVA with Dunnet’s multiple comparisons correction, F (1.001, 9.009) = 283.8. (c) Model of the lipid A modifications in colistin-resistant cells, in which the *arn* operon is activated by PhoPQ, resulting in 4-amino-4-deoxy-ʟ-arabinose modification of lipid A and colistin resistance. (d-e) MALDI-TOF-MS spectra of strain RS following growth in (d) broth alone or (e) or 100 μg/mL colistin, a single spectra representative of 3 biological replicates is shown. (f) PAP of strain RS wildtype, Δ*phoPQ*, and Δ*arnB* mutants on Mueller-Hinton agar (MHA) containing colistin; from three independent experiments with n=9 total biological replicates for RS and Δ*arnB*, n=12 total biological replicates for Δ*phoPQ*. (g) P*_arn_gfp* expression intensity in strain RS wildtype, from three independent experiments; data were determined from 700-1000 cells for each experiment.

While the majority susceptible population of cells did not have detectable ւ-Ara4N modifications to lipid A (Figure 1d, Supp. Table 2), which can confer colistin resistance, these modifications were prominent after the resistant subpopulation was enriched in broth containing colistin (Figure 1e, Supp. Table 2). Correspondingly, genetic deletion of *arnB* which encodes UDP-4-amino-4-deoxy-ʟ-arabinose-oxoglutarate aminotransferase, converts UDP-4-ketopentose to UDP-ʟ-Ara4N, and is required for ւ-Ara4N addition to lipid A (*22*), resulted in loss of the colistin resistant subpopulation (Figure 1f). Similarly, deletion of *phoPQ*, encoding the two-component system that activates expression of the *arn* operon (*16, 17*), resulted in elimination of the highly resistant subpopulation (Figure 1f, Supp. Figure 3a-b). Further, mutants lacking *arnB* or *phoQ* exhibited no ւ-Ara4N modifications to lipid A (Supp. Figure 3c-d). Thus, the colistin resistant subpopulation is dependent on PhoPQ and ւ-Ara4N modification of lipid A.

### Activation of the *arn* operon promoter is not bimodal

Since the *arn* genes are critical for the colistin resistant subpopulation, we hypothesized that heterogeneous activation of the *arn* promoter might generate the phenotypic heterogeneity in colistin HR in strain RS. If correct, we predicted ∼1% of the population would have high *arn* expression and demonstrate resistance, while ∼99% of the cells would have low *arn* expression and be susceptible. To test this hypothesis, we generated a chromosomally-encoded transcriptional reporter by fusing the *arn* promoter to *gfp* (P*_arn_gfp*). We imaged RS::P*_arn_gfp* and found that, counter to our hypothesis, P*_arn_gfp* expression was not bimodal but rather unimodal with normal distribution (Figure 1g). These data demonstrate that while *arn* expression is required for colistin HR, its expression is not sufficient to generate population heterogeneity.

### Activation of the alternative sigma factor σ^E^ pathway increases colistin resistance

To attempt to identify the factors that generate the phenotypic heterogeneity in colistin HR, we first performed a transposon screen for mutants that do not survive on colistin at wildtype levels. Of ∼16,000 colonies screened, we identified 42 transposon mutants that demonstrated reduced frequency of the colistin resistant subpopulation. Validating our screen, we identified a mutant in *arnD*, part of the *arnBCADTEF* operon that is required for colistin resistance (Supp. Table 3). We noticed that transposon mutants were identified in *clpP*, which encodes a protease adaptor involved in many cellular pathways, and *sspB*, encoding the stringent starvation protein B, which acts in conjunction with ClpP in the activation of the alternative sigma factor σ^E^ (encoded by *rpoE*) (Figure 2a). We therefore pursued the hypothesis that σ^E^ may contribute to colistin HR. Clean deletion mutants of either *clpP* or *sspB* displayed increased susceptibility to colistin, further confirming the accuracy of the transposon screen, although the degree of susceptibility of the *clpP* mutant was much greater than that of *sspB* (Figure 2b, Supp. Figure 4a). An *rpoE* mutation was not identified in our screen, and we were unable to generate an *rpoE* deletion, consistent with its reported essentiality in *E. coli* (*23*). To further test the involvement of σ^E^, we instead deleted *rseA*, encoding its anti-sigma factor which restrains its activity, and *rseB*, which encodes a periplasmic protein that enhances RseA inhibition of σ^E^ (*24, 25*). Inactivation of RseA, and to a lesser extent RseB, is known to increase σ^E^ activity (*24, 25*). The frequency of the colistin resistant subpopulation increased in both the Δ*rseA* and Δ*rseB* mutants relative to wildtype (Figure 2c). In the case of the Δ*rseA* mutant, the frequency of the resistant cells reached almost 100% even when exposed to 100 μg/mL colistin (Figure 2c, Supp. Figure 4b-c). Taken together, the genetic inactivation of proteins which promote σ^E^ activation (Δ*clpP*, Δ*sspB*) resulted in a lower frequency of the colistin resistant population, and inactivation of proteins which restrain σ^E^ activation (Δ*rseA*, Δ*rseB*) increased the frequency of cells resistant to colistin. These results demonstrate the role of the σ^E^ pathway in colistin HR.

**Figure 2.**
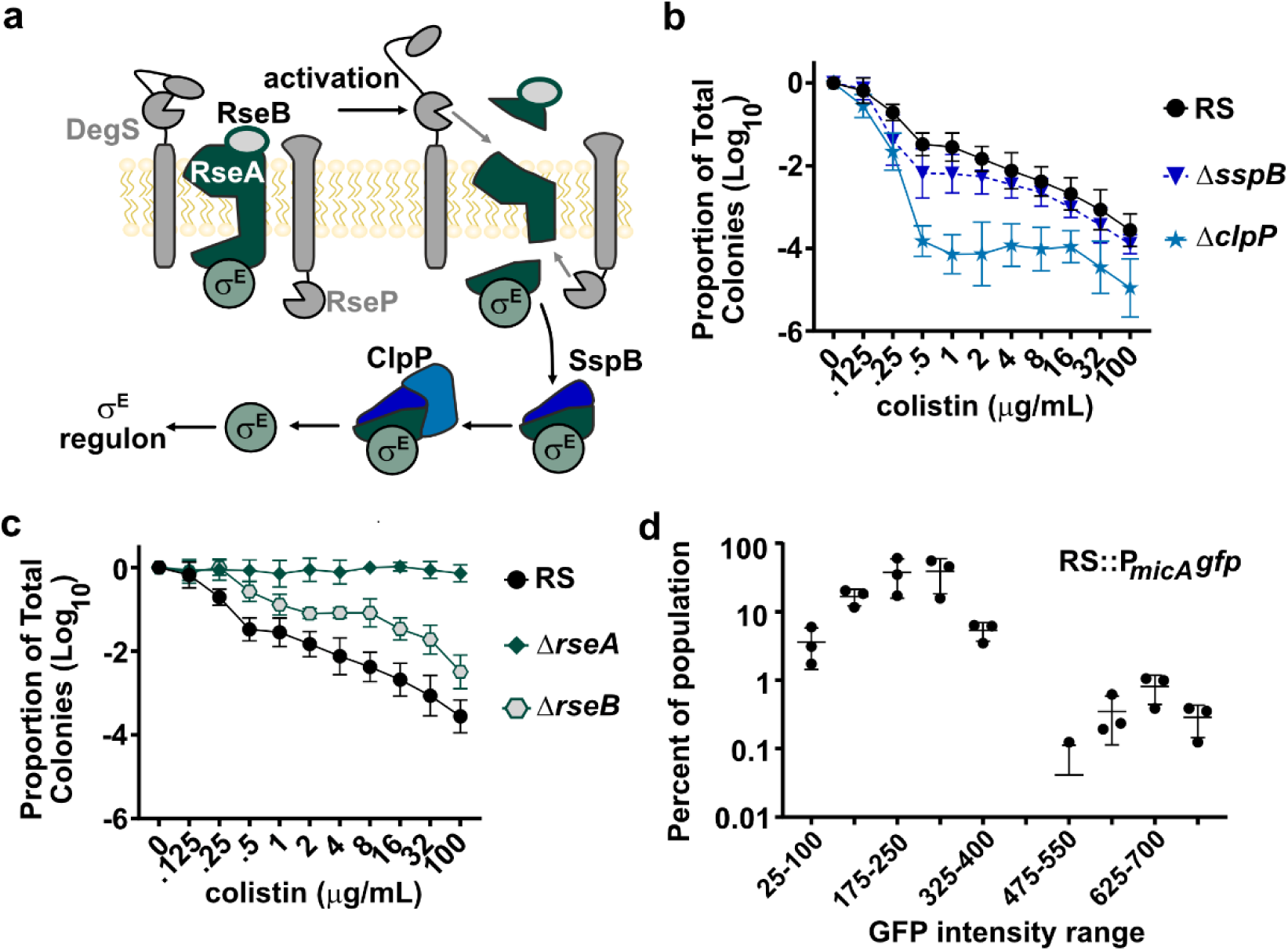
Activation of alternative sigma factor σ^E^ pathway increases colistin resistance. (a) Model of σ^E^ activation: following proteolytic degradation of its anti-sigma factor RseA and accessory protein RseB by DegS and RseP, σ^E^ is liberated from the membrane. The protease ClpP with the adaptor protein SspB degrade the remaining RseA domain to allow σ^E^ to transcriptionally regulate its large regulon. (b) Population analysis profile (PAP) of strain RS and Δ*sspB* and Δ*clpP* mutants. (c) PAP of strain RS and Δ*rseA* and Δ*rseB* mutants; (b-c) are from the same two independent experiments with n=6 total biological replicates for RS and Δ*rseB* and n=8 total biological replicates for the rest of the strains. (d) The *micA* promoter (σ^E^-controlled) was used to monitor σ^E^ activity: P*_micA_gfp* expression intensity in wildtype strain RS, from three independent experiments; data were determined from 700-1000 cells for each experiment.

### σ^E^ promoter activation is bimodal

Having identified the σ^E^ pathway as an important regulator of colistin HR, we returned to our hypothesis that bimodality in gene expression could generate HR. We therefore generated a chromosomally-encoded reporter of σ^E^ activity using the promoter for a σ^E^-regulated gene, *micA* (RS::P*_micA_gfp*) (*26*). We then microscopically imaged individual cells of the RS::P*_micA_gfp* strain and measured their fluorescence. We observed that σ^E^ activity demonstrated bimodality, in which ∼1% of the population showed high P*_micA_gfp* expression (Figure 2d), which matches the measured percentage of colistin resistant cells when RS is grown in the absence of colistin (Figure 1a). Together, these data suggest that active σ^E^ in a rare subpopulation of cells could generate the colistin resistant subpopulation.

### Rare σ^E^ high cells constitute the colistin resistant subpopulation

To investigate the relationship between σ^E^ activity and survival on colistin, we next assessed the P*_micA_gfp* σ^E^ reporter strain before (Figure 3a) and after addition of 8 μg/mL colistin (Figure 3b, c) and performed single-cell, time-lapse microscopy to track the fate of each cell upon colistin exposure (Supplemental Video). We designated each cell as susceptible or resistant based on its ability to replicate in the presence of colistin. Intriguingly, cells that did not express the P*_micA_gfp* reporter did not replicate in the presence of colistin, whereas only cells with high reporter expression prior to colistin exposure were able to grow and divide (Figure 3d). Further, when we encoded the P*_micA_gfp* reporter in the Δ*rseA* strain, which is made up almost exclusively of colistin resistant cells (Figure 2c), we observed a corresponding and robust increase in P*_micA_gfp* expression as compared to the parental wild-type RS strain (Figure 3e). Together, these data suggest that the resistant subpopulation in RS is generated by pre-existing, bimodal σ^E^ activity prior to colistin exposure.

**Figure 3.**
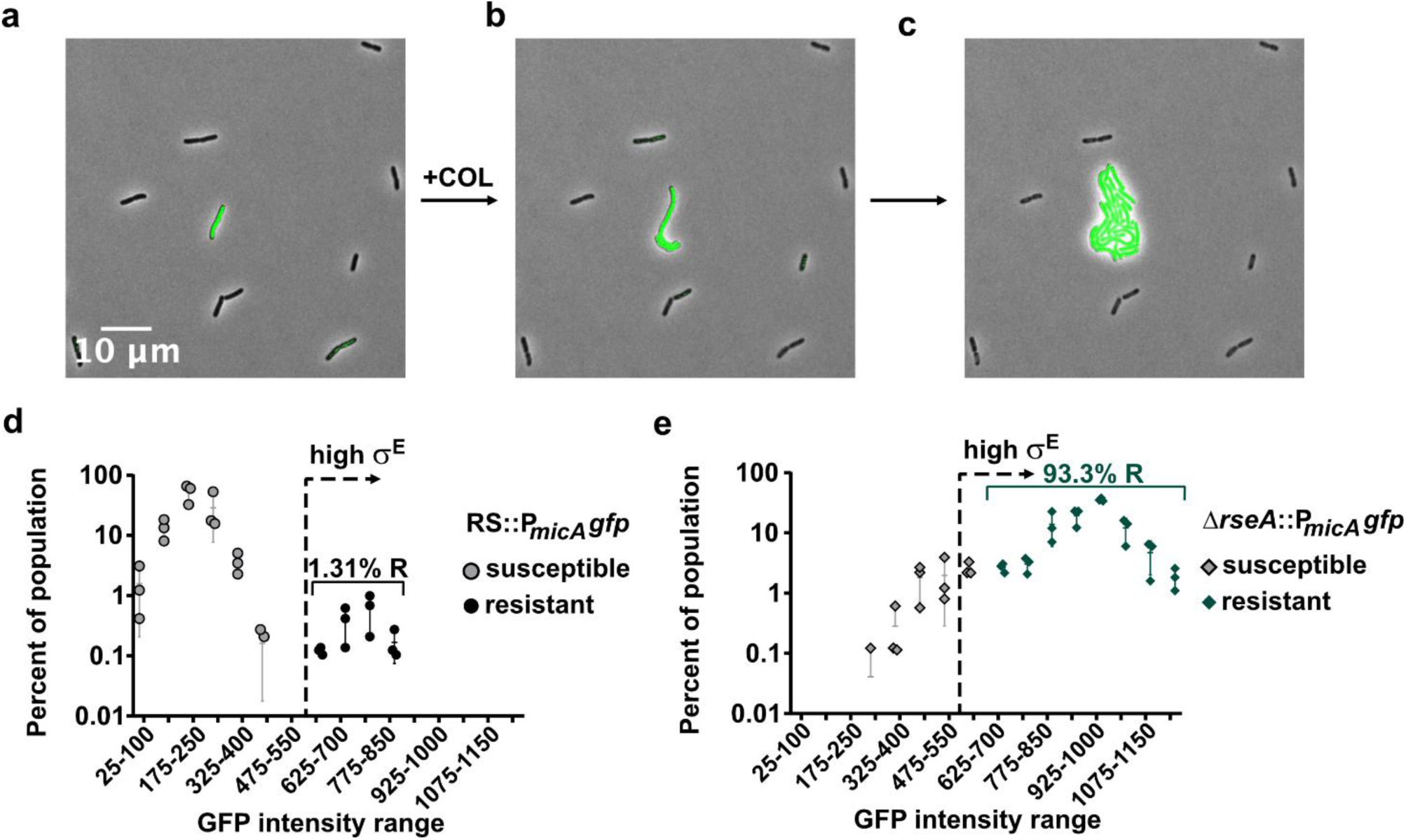
σ^E^ promoter activation is bimodal and predictive of colistin resistance. (a-c) Representative images of P*_micA_gfp* expression (a) prior to exposure to 8 μg/mL colistin, (b) after 2 h of colistin exposure and (c) after 4 h of exposure. (d-e) P*_micA_gfp* expression intensity in strain (d) RS wildtype or (e) the Δ*rseA* mutant prior to colistin exposure. The cell fate was monitored over time after adding 8 µg/mL colistin, and each cell was designated as susceptible or resistant. (d-e) are from three independent experiments; data were determined from 700-1000 cells for each experiment.

### Colistin heteroresistance conferred by σ^E^ activation requires ւ-Ara4N modification

It was unclear how activation of the σ^E^ pathway led to colistin HR and if lipid A modifications were required. We performed PAP of a strain lacking both *rseA* and *arnB* and observed that deletion of *arnB* rendered the highly resistant Δ*rseA* strain susceptible to colistin (Figure 4a, Supp. Figure 5). Next, we directly characterized the lipid A of the Δ*rseA* and Δ*arnB rseA* strains grown in the absence of colistin. Compared to wildtype RS, the Δ*rseA* strain demonstrated extensive modifications with ւ-Ara4N, consistent with its high frequency of colistin resistant cells. However, these modifications were absent in the Δ*arnB rseA* strain (Figure 4b). Thus, the colistin resistance conferred by constitutive activation of σ^E^ requires *arn*-mediated lipid A modification. Since *arn* plays a critical role in lipid A modification and colistin resistance, we examined its expression at the single-cell level and its relationship with survival on colistin using the P*_arn_gfp* reporter in wildtype RS. While *arn* expression overall was unimodal (Figure 1g), we observed a trend where the cells with higher *arn* expression were more likely to survive on colistin (Figure 4c), although there was overlap in *arn* expression between the resistant and susceptible cells. This was in contrast to the results for *micA* expression which was bimodal and where there was no overlap between the resistant and susceptible populations.

**Figure 4.**
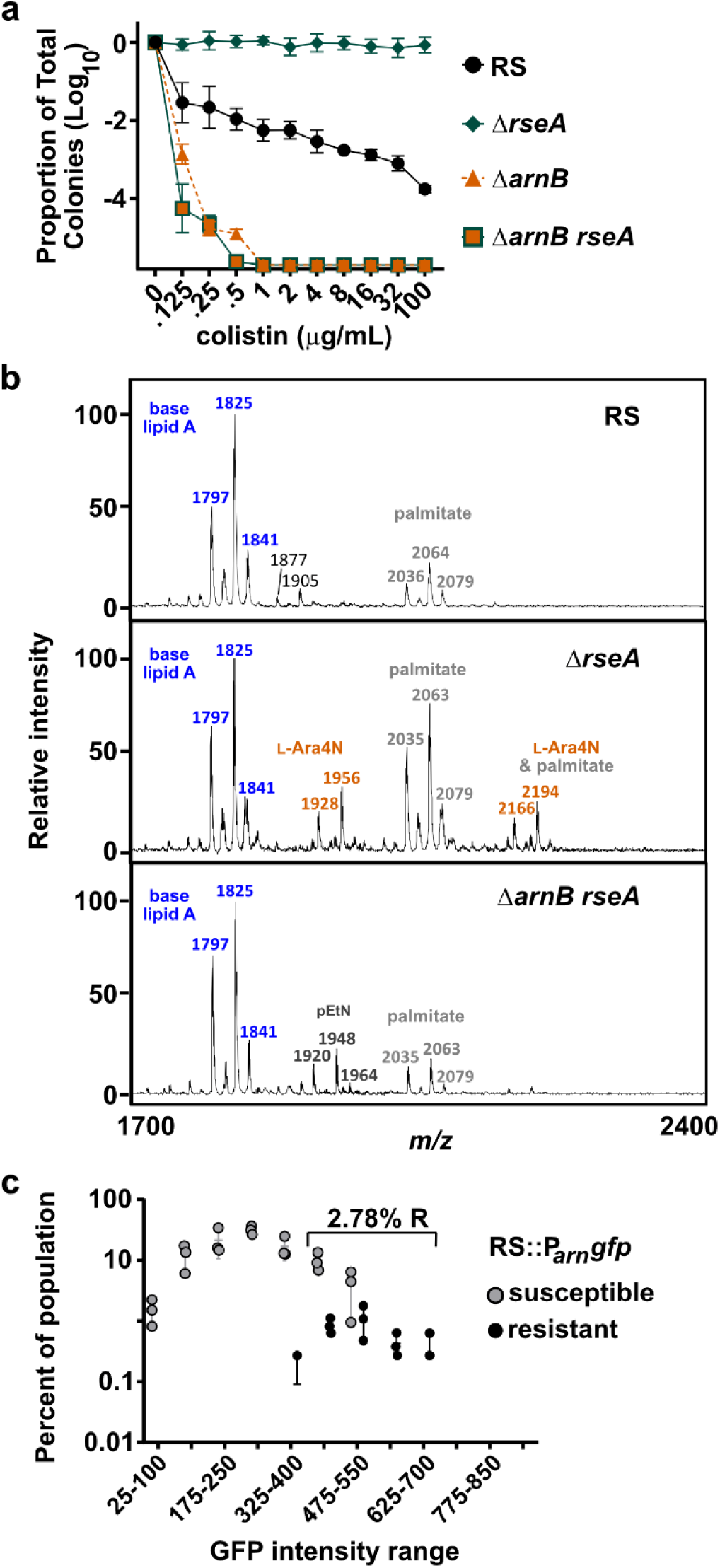
σ^E^ confers colistin resistance by increasing *arn* levels. (a) Population analysis profile (PAP) of strain RS wildtype, and the Δ*rseA*, Δ*arnB*, and Δ*arnB rseA* mutants, from two independent experiments with n=7 total biological replicates. (b) MALDI-TOF-MS spectra of strain RS, Δ*rseA,* and Δ*arnB rseA* following growth in broth alone; a single spectra representative of 3 biological replicates is shown. Spectra of RS at top of B is reproduced from Figure 1d for the purpose of comparison. (c) P*_arn_gfp* expression intensity in strain RS wildtype or prior to colistin exposure. The cell fate was monitored over time after adding 8 µg/mL colistin, and each cell was designated as susceptible or resistant, from three independent experiments; data were determined from 700-1000 cells for each experiment.

### The σ^E^ pathway generates heterogeneity but is not sufficient for heteroresistance

PhoPQ is involved in *arn* induction and colistin HR in RS, yet it was not clear if or how its activity impacted σ^E^ activation. The heterogeneous activation of σ^E^, as measured by the *micA* reporter, was maintained in the *phoPQ* mutant (Figure 5a, compared to Figure 2d), indicating that PhoPQ is not required for this phenomenon. This finding is also supported by the PAP of the *phoPQ* mutant, where a subpopulation of roughly 1 in 100,000 cells with a higher MIC than the main population was present at 0.25, 0.5, and 1 ug/ml colistin, demonstrating heterogeneity (Figure 1f). However, none of the cells in the *phoPQ* mutant were classified as colistin resistant, since none of them survived at the clinical breakpoint concentration of 4 μg/ml. This is consistent with vastly reduced *arn* reporter expression in the *phoPQ* mutant (Figure 5b, compared to Figure 1g). This highlights that 1) overall population phenotypic heterogeneity (heterogeneous survival) and heteroresistance (resistance of a subpopulation at the clinical breakpoint) are separable traits in strain RS, and 2) that PhoPQ is not required for phenotypic heterogeneity, but is required for the overall increase in MIC of a subpopulation to and beyond the clinical breakpoint, thus making RS heteroresistant to colistin. These results demonstrate that the σ^E^ pathway is the generator of phenotypic heterogeneity in RS and PhoPQ is a generator of resistance, with the combined activity of these two regulatory systems being required to generate colistin heteroresistance.

**Figure 5.**
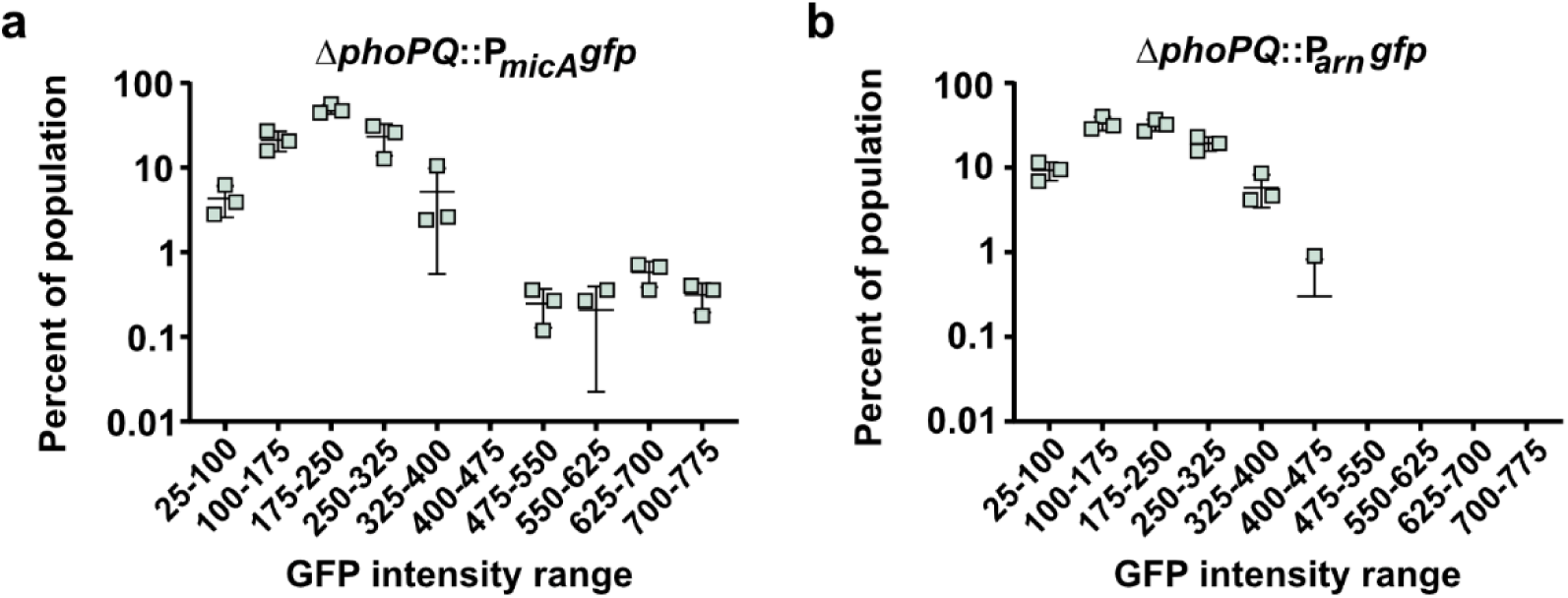
PhoPQ does not affect σ^E^ pathway heterogeneity. (a) P*_micA_gfp* expression intensity and (b) P*_arn_gfp* expression intensity in strain RS Δ*phoPQ*, from three independent experiments; data were determined from 700-1000 cells for each experiment.

## DISCUSSION

Clinical isolates are conventionally categorized as susceptible or resistant to a given antibiotic, classifications which omit the phenotypic heterogeneity observed in heteroresistance. Colistin HR is a frequent feature of clinical *Enterobacter* isolates, is often undetected by conventional antibiotic susceptibility testing, and can cause treatment failure in murine models of infection. However, it was still unclear what generated the phenotypic heterogeneity in this system.

Here, we performed a transposon screen and identified a critical role for the alternative sigma factor σ^E^ in colistin HR (Figure 2). In a population of wildtype cells of strain RS, prior to colistin exposure, ∼1% of the population exhibited increased σ^E^ activity, making up a distinct second peak in the bimodal distribution of σ^E^ activity (Figure 2d). Single-cell tracking revealed that exclusively these cells with high σ^E^ activity survived colistin exposure (Figure 3d). Although PhoPQ has been strongly linked to colistin HR, we were surprised to discover that this system is not the generator of phenotypic heterogeneity, which was maintained in a *phoPQ* mutant strain (Figure 5a). However, the heterogeneity was lost, as evidenced by a unimodal curve of σ^E^ activity, when σ^E^ was constitutively activated via deletion of the σ^E^ inhibitory anti-sigma factor, *rseA* (Figure 3e), further demonstrating that the σ^E^ pathway is the generator of heterogeneity. While phenotypic heterogeneity was dependent on the activity of the σ^E^ pathway, HR to colistin nonetheless required PhoPQ as none of the cells of the *phoPQ* mutant survived at the colistin clinical breakpoint (Figure 1f). Importantly, the cells surviving colistin exposure also had higher *arn* expression (Figure 4c), which required PhoPQ (Figure 5b). These data highlight that PhoPQ is not required for phenotypic heterogeneity, but is required for the overall increase in MIC of a subpopulation to the clinical breakpoint, thus generating colistin HR. Interestingly, these data demonstrate that overall population phenotypic heterogeneity (heterogeneous survival) and heteroresistance (resistance of a subpopulation at the clinical breakpoint) are separable traits in strain RS. Further, these results reveal that the σ^E^ pathway is the generator of phenotypic heterogeneity in RS, and PhoPQ is a generator of resistance, with the combined activity of these two regulatory systems being required to generate colistin heteroresistance. Together, these data elucidate a novel mechanistic paradigm for heteroresistance via the combinatorial activity of distinct regulatory pathways (Figure 6).

**Figure 6.**
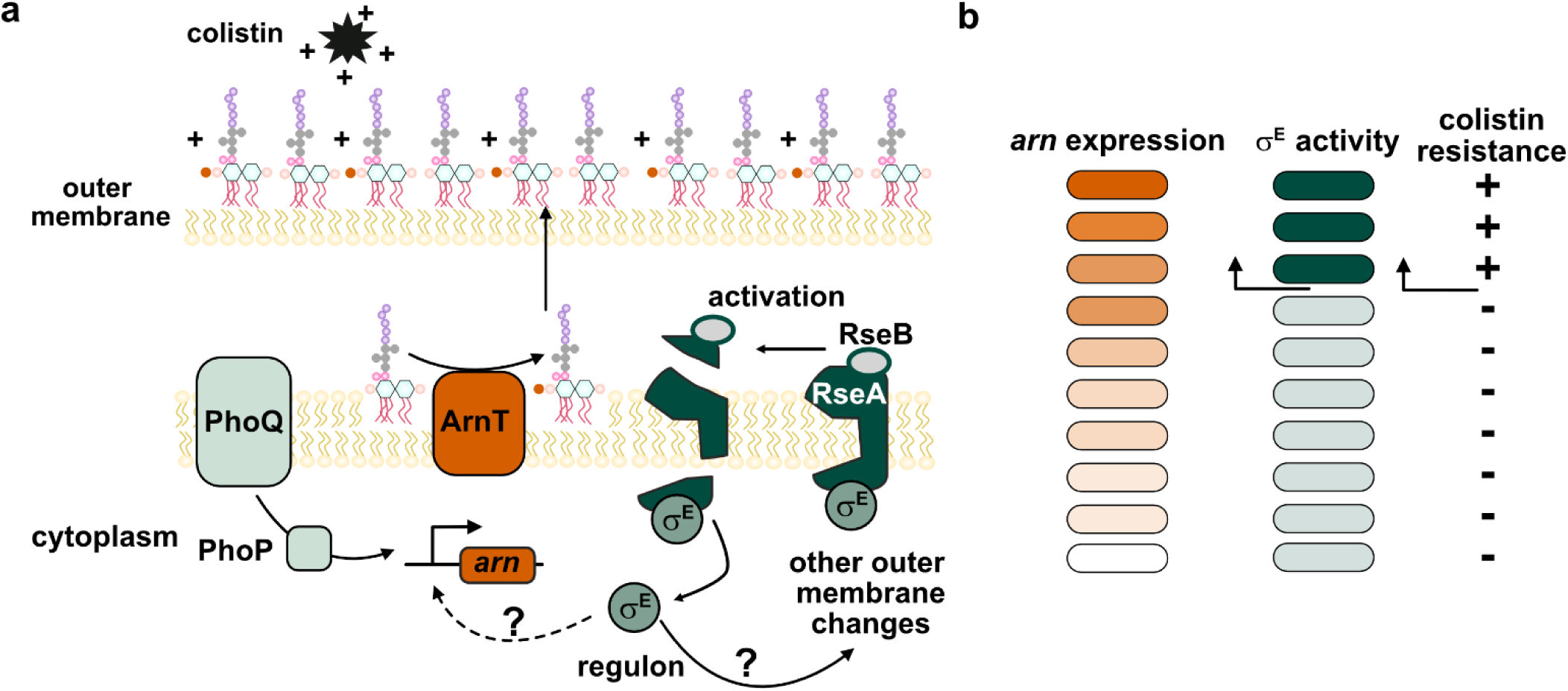
Proposed model for bimodality in σ^E^ generating a colistin-resistant subpopulation. (a) A proposed model for the activity of signaling pathways active in the subpopulation of cells with pre-existing colistin resistance. PhoPQ directly activates transcription of the *arn* operon, while activation of σE additionally directly or indirectly increases *arn* transcription, resulting in lipid A modification that prevents colistin entry. (b) Prior to colistin exposure, *arn* promoter activity follows a broad unimodal expression, while σ^E^ promoter activity is distinctly bimodal. High σ^E^ cells have higher *arn* promoter activation and are colistin-resistant, while cells with moderately high P*arn* activity but low σ^E^ activation remain colistin-susceptible.

This combinatorial regulatory model of HR is quite distinct from the best understood mechanism of HR in which one event, an increase in copy number of a given single gene, generates HR. The clearest and most commonly observed example thus far has been the tandem gene amplification of a DNA segment including a beta-lactamase gene and leading to overproduction of the beta-lactamase in the cells in the population with increased gene copy number. This mechanism can generate a broad continuum of cellular subpopulations resistant to a given beta-lactam antibiotic (*27*). It should be noted that gene amplification is capable of causing colistin HR as this was observed in one instance in *S.* Typhimurium (*28*), although this does not appear to be a common mechanism at least among *Enterobacter* isolates.

Interestingly, using the *rseA* mutant in which σ^E^ activity is constitutive, we observed that the σ^E^ pathway can also impact *arn* expression. In the *rseA* mutant, the entire population demonstrated a shift to increased *arn* expression (Supp. Figure 6g) and colistin resistance (Figure 2c). However, *phoPQ* deletion did not significantly reduce *arn* expression in the *rseA* mutant (Supp. Figure 6c, Supp. Figure 7) and only led to a modest reduction in the frequency of the resistant subpopulation to ∼1 in 10 cells (Supp. Figure 6a). This was in contrast to the phenotype of the *rseA/arnB* double mutant where no resistant cells remained (Figure 4a). This highlights that in the setting of robust, constitutive σ^E^ activation, *arn* plays a more critical role in colistin HR than PhoPQ, and that σ^E^ activation can induce *arn*.

Since many host antimicrobial peptides are highly cationic, similar to colistin, we hypothesized that antimicrobial peptide treatment would activate the σ^E^ pathway. When strain RS was treated with the human antimicrobial peptide LL-37, we observed an increase in transcript abundance of *rpoE* and *rseA*, consistent with activation of the σ^E^ pathway (Supp. Figure 8). These conditions also activated the canonical PhoPQ response to LL-37, as measured by expression of the PhoPQ-activated *mgtA* transcript and *arnB* (Supp. Figure 8a). Consistent with a σ^E^ pathway response to LL-37, the Δ*rseA* strain was more resistant to LL-37 killing than wildtype (Supp. Figure 8b). Thus, these data demonstrate that σ^E^ is involved not only to survival in colistin, but also in response to host antimicrobial peptides.

It is not clear why lipid A is not modified by ʟ-Ara4N in every cell, constitutively., leading to homogeneous resistance to colistin and other CAMPs. There is a fitness cost associated with constitutive σ^E^ activation, which is toxic in *E. coli* (reviewed in (*21*). Thus, HR may serve as a form of bet-hedging, in which a dynamic pre-existing subpopulation can co-exist with other phenotypically diverse subpopulations, and can be transiently enriched when the selective pressure is present. The mammalian host environment is one such environment, as the colistin resistant subpopulation in strain RS is enriched during murine infection by the selective pressures of lysozyme, NADPH oxidase, and the antimicrobial peptide mCRAMP, even in the absence of colistin treatment (*16*). Thus, bimodality in σ^E^ activation may confer on the population an advantage in resistance to antibiotic and host antimicrobial stresses, without a commitment of the entire population to costly LPS modifications.

Our model of combinatorial regulatory signaling converging on the *arn* resistance determinant is consistent with work on colistin HR in other *Enterobacter*, where inactivating PhoPQ/*arn* eliminates the resistant subpopulation, and manipulating PhoPQ activity changes the frequency of the resistant subpopulation, but not the baseline heterogeneity in the population (*8, 17*). In sum, we have identified a transcriptional regulator that generates phenotypic heterogeneity to colistin in the population. This heterogeneity facilitates HR when the PhoPQ two-component system increases the overall resistance level of all the cells in the population. Thus, this combinatorial regulatory model is a new paradigm in the mechanistic generation of HR, a phenomenon which is increasingly recognized as a cause of antibiotic treatment failure in the clinic.

## ACKNOWLEDGMENTS

pBAV1K-T5-gfp was a gift from Ichiro Matsumura (Addgene plasmid # 26702; http://n2t.net/addgene:26702; RRID:Addgene_26702). We thank Edgar Sherman for generating the Δ*arnB rseA* strain.

## Author contributions

Conceptualization: JEC, EKC, MM, MK, DSW. Investigation: JEC, EKC, MM, LMV, MKK. Visualization: JEC, MM, LMV. Funding acquisition: MK, RKE, DSW. Supervision: MK, RKE, DSW. Writing-original draft: JEC and DSW. Writing-review and editing: all authors

## Data and materials availability

all data needed to evaluate the conclusions in the paper are present in the paper and/or the Supplementary Materials. Source data is provided as a Supplementary File.

## MATERIALS AND METHODS

### Statistical analysis and data presentation

The number of experiments and number of biological replicates are detailed in the figure legends. PAP graphs show the mean and standard deviation; alternative graphs showing individual data points are in Supplemental Figures. Other graphs show each data point and mean, and error bars indicate standard deviation as applicable. Violin plots show median and quartiles. Statistical analysis was performed in Graphphad Prism version 10.4.2 and details of each test are in figure legends.

### Isolate information

*Enterobacter cloacae* complex strain RS was isolated from a blood sample from a renal transplant recipient at Emory University Hospital, Atlanta, GA and described previously (*16, 29*). The RS genome is available in NCBI Genbank, accession numbers CP010512 and CP010513. *E. cloacae* ColS1 and ColR2 are clinical isolates and described previously (*29*). Strains and plasmids in this paper are detailed in Supp. Table 4.

### Reagents

Mueller-Hinton agar (MHA; BD Difco) and Mueller-Hinton broth (MHB; BD Difco) were used throughout as indicated. For growth of *E. cloacae* RS before plating on MHA containing colistin, bacteria were grown at 37°C with shaking, 1.5 mL volume in 12x75 mm polypropylene aeration tubes (Globe Scientific #110438). YESCA was made by autoclaving 1 g/L yeast extract (Fisher BP1422) and 10 g/L casamino acids (Acros #61204) in water. YESCA agar included 20 g/L agar (Sigma A1296).

Antibiotics used were: colistin sulfate (Sigma #C4461), kanamycin sulfate (Teknova #K2150; 90 µg/mL for strain RS), tetracycline (Alfa Aesar #J61714; 20 µg/mL for RS), chloramphenicol (Fisher #BP904, 50 µg/mL for RS) and spectinomycin dihydrochloride pentahydrate (Sigma #S4014; 50 µg/mL for *E. coli*, 300 µg/mL for RS).

Medium gel pads used for cell imaging under microscope: UltraPure^TM^ agarose (Invitrogen #16500100) and MHB was used to prepare MHB 1.5% agarose gel pad.

### Population analysis profile

Population analysis profile (PAP) was performed as indicated in Supp. Figure 1. A given strain was grown overnight from a single colony streaked to MHA from -80°C glycerol stocks. After approximately 16-20 h of growth, the culture was diluted 1:1000 (except for strains with *rseA* or *clpP* mutation, which were diluted 1:250 to account for diminished growth yield of these strains) into fresh media and grown until OD reached ∼0.3. The cultures were serially diluted in PBS in a 96-well plate (Falcon) and 7.5 or 10 µl of each dilution was plated on MHA containing colistin as indicated. Colonies were enumerated after 24-48 h of growth. For many experiments shown, the same protocol was followed but the cultures were plated only on MHA containing 0 or a single colistin concentration indicated.

The surviving colonies are enumerated and the isolate is classified as resistant if at least 50% of the total colonies grow at 1 or 2X breakpoint. An isolate is considered susceptible if less than 0.0001% (−6 logs) of the cells grow at any concentration shown. An isolate is considered heteroresistant if there is greater than 0.0001% survival at 1X and 2X breakpoint and an ≥8-fold difference between the resistance of the subpopulation and resistance of the main population. As an example for RS, <50% of the population survives at 0.125 µg/mL, thus the MIC_susceptible population_ is 0.125 µg/mL. The subpopulation survives up to 100 µg/mL, so the MIC_resistant population_ of the resistant population is > 100 µg/mL. For RS the fold change of the MIC_resistant population_ relative to MIC_susceptible population_ is >8. The limit of detection is approximately -5.5-6 logs but varies based on the density of the culture, in the figures the y-axis is set to the approximate limit of detection for each graph.

In defining the features of heteroresistance in the isolates of this work, based on guidelines set forth by the field (*6, 9, 30*):

1. Clonality: these isolates demonstrate monoclonal heteroresistance, they are purified isolates and single colonies are used throughout experiments.
2. Level of resistance: the MIC of the resistant subpopulation in wildtype RS is ≥8X the MIC of the main population, when comparing the amount of killing by the lowest concentration of colistin used throughout, and the growth of the resistant subpopulation at 100 µg/mL.
3. Frequency of the resistant subpopulation: We consider the frequency of the subpopulation at 2X the CLSI breakpoint, for wildtype RS the frequency is ∼0.1-1%.
4. Instability in the frequency of the resistant subpopulation: wildtype RS demonstrated unstable heteroresistance. As shown in the passage experiments after selection, there was a significant reduction in the resistant population frequency within a single passage (∼10 generations per passage) in antibiotic free media.

### Copy number variation sequencing analysis

Illumina sequencing of samples of RS previously grown in Mueller Hinton broth containing 0 or 100 μg/mL colistin (*16*) were accessed from NCBI SRA. Reads were mapped to the respective reference genome with CNV analysis. Quality control and adapter trimming was performed with bcl2fastq(*31*). Reads were mapped to their respective references via bwa mem(*32*). PCR and optical duplicates were marked and excluded from the analysis using PicardTools’ ‘MarkDuplicates’(*33*) functionality. Aligned read counts were imported into R’s CNOGpro(*34*)package. CNV events were called via CNOGpro using a bootstrapping method, which calculates an average gene number event giving a possible upper and lower bound. Genome sequencing and analysis were performed by SeqCenter (https://seqcenter.com). Supplemental Table 1 details analysis results as well as SRA accessions for each sample.

### Resistance stability assays

RS was streaked from -80°C glycerol stocks to MHA for isolation and a single colony was used to start overnight cultures in 1.5 mL of MHB and grown 15-20 h. 15 µl of the cultures were diluted into 1.5 mL of MHB and grown until OD reached ∼0.3-0.4. These cultures were diluted and plated MHA containing 0 or 100 µg/mL colistin, the baseline. 15 µL of the cultures were added to 1.5 mL fresh MHB containing 100 µg/mL colistin and grown for 20 h. Following growth, the cultures were diluted and plated MHA containing 0 or 100 µg/mL colistin. 1.5 µL of the cultures were added to 1.5 mL fresh MHB and grown until OD reached ∼0.3-0.4. Following growth, the cultures were diluted and plated MHA containing 0 or 100 µg/mL colistin.

### Lipid A mass spectrometry

Sample growth: strain RS and indicated mutants were grown overnight in 1.5 mL MHB and 120 µL was added to 12 mL fresh MHB in a 50 mL conical tube and grown for 4 h, then harvested by centrifugation. For RS grown in colistin, 100 µL of overnight culture was added to 10 mL MHB containing 100 µg/mL colistin in a 50 mL conical tube and grown for 9 h, then harvested by centrifugation. Cells were washed in PBS and stored in -80°C.

Fast lipid analysis technique (FLAT)(*35*) was adapted as follows: bacteria were scraped from frozen pellets and spotted directly onto a steel MALDI plate. 1uL of FLAT extraction buffer (0.2 M anhydrous citric acid, 0.1 M trisodium citrate dihydrate) was spotted on top of the plated bacteria. The MALDI plate was placed into a humidified 100 °C “panini-press” chamber for 30 minutes. Afterwards, the plate was rinsed with endotoxin-free water and air dried. 1uL of Norharmane matrix (10 mg/mL in 2:1 (v/v) chloroform-methanol solution) was spotted on each sample. Samples were analyzed in linear, negative ion mode on a Bruker Microflex LRF. Data was processed with flexAnalysis software.

### Imaging of cells and GFP expression

RS wildtype or mutants encoding *att*:: P*_arn_gfp+* or *att*:: P*_micA_gfp+* were used to visualize population-level variation in promoter activation prior to colistin exposure, and to assess viability following treatment. Imaging of cells and GFP expression was performed as previously described (*36, 37*). When a cell culture reached an OD_600_ of approximately 0.1, cells were transferred onto a 35 mm glass-bottom Petri dish (Cellvis) and overlaid with Mueller-Hinton Broth (MHB) gel pad containing 1.5% agarose. Cells were imaged using an Olympus IX83 inverted microscope with a 60× oil immersion phase-contrast objective, housed within a temperature-controlled (37 °C) incubation chamber (InVivo Scientific). GFP fluorescence was detected using a GFP filter cube. Snapshot imaging of cells was performed before exposure to colistin. Time-lapse images of cells were captured at 30-minute intervals after adding colistin to the agarose pad covering cells.

### Image analysis

Fluorescence intensity of GFP was measured using a plug-in for Fiji ImageJ software, named MicrobeJ (*38*) (version 5.13 I (22)). Reported fluorescence intensities represent intracellular GFP signals after subtracting both extracellular background and the autofluorescence of wildtype RS cells. Cell viability under colistin treatment was determined through time-lapse imaging. Continuous cellular division and colony formation represented survival. Killing of cells by colistin was defined as visible lysis or permanent arrest of cell growth.

### Transposon screen

750 µl of a culture of a RS *att*::P*_arn_gfp*+ was combined with 750 µl *E. coli* BW19851 carrying pFD1, collected by centrifugation and resuspended in 50 µl YESCA (yeast extract casamino acids broth). The suspension was plated on a 0.45 µM nitrocellulose membrane placed on top of a YESCA agar plate and incubated at 37°C for 2.5 h. The nitrocellulose was washed in a 15 ml tube with YESCA containing 1 mM IPTG and 300 µg/mL spectinomycin, grown for 3 h at 37°C, diluted and plated on MHA containing 300 µg/mL spectinomycin and 90 µg/mL kanamycin. After 24 h of growth at 37°C, colonies were replica-plated using sterile velvet onto MHA containing kanamycin or kanamycin and 100 µg/mL colistin. Colonies that appeared on kanamycin but not kanamycin+colistin were restreaked for isolation. To identify the transposon insertion site, colony PCR was performed using primers 71/72 to yield the DNA flanking the transposase. 2 µl of the PCR product from JCP71/72 was the template for subsequent nested PCR with JCP73/74. PCR products were Sanger sequenced (Azenta) using JCP74 and aligned to the RS genome.

### Strain construction: gene deletions

Lambda-red based allelic exchange was used to replace the coding sequence of genes with a kanamycin resistance gene as indicated. The kanamycin resistance gene from pEXR6K_kanFRT was cloned using Promega GoTaq 2X master mix with Flp recognition sequence and homology to regions flanking the gene to be replaced using primers JCP171/172 (*phoPQ*), EKC202/203 (*arnB*), EKC301/302 (*sspB*) JCP59/60 (*clpP*) EKC309/310 (*rseA*) and JCP56/57 (*rseB*), detailed in Supp. Table 5. The purified PCR product was electroporated into competent RS (or mutant) carrying pKD46-tet and transformants were selected on kanamycin. Transformants were re-streaked to kanamycin for isolation and subject to PCR for successful allelic exchange as indicated by a product size change using primers flanking the gene to be replaced: primers JCP173/174 (*phoPQ*), EKC208/209 (*arnB*), EKC303/304 (*sspB*) JCP61/62 (*clpP*) EKC311/312 (*rseA*) and JCP58/EKC311 (*rseB*) detailed in Supp. Table 5. As indicated, the kanamycin resistance gene was removed with Flp recombinase(*39, 40*) to create unmarked in-frame deletions The mutants were then made electrocompetent and electroporated with pCP20(*40*). Transformants were selected at 30°C on chloramphenicol, patched to chloramphenicol at 30°C and grown for 24 h, then patched to MHA and MHA+kanamycin. Kanamycin-sensitive mutants were streaked for isolation and PCR was used with the same flanking primers to screen for the loss of the kanamycin resistance gene.

### Strain construction: chromosomally integrated plasmids

For P*_rmicA_gfp*, pCD13PSK was digested with BamHI, the *micA* promoter region was an ultramer JCP41, and promoterless *gfp+* was cloned with JCP31/42 from the vector pZEP07 (*41*) as template. For P*_arn_gfp,* pCD13PSK *gfp*+ was amplified with JCP556/557 and the *arn* promoter cloned with JCP558/559.

For pCD13PSK *phoPQ*, P*_phoP_phoPQ* was cloned with EKC223/235 and assembled with pCD13PSK digested with XhoI and XbaI. For pCD13PSK *arnBCADTEF*, the entire arn operon with native promoter was cloned with JCP67/68 and assembled with pCD13PSK digested with BamHI and SacI. For each construct, PCR and plasmid components were assembled with NEB Hifi assembly according to manufacturer’s instructions and transformed. Plasmids were propagated in *E. coli* PIR2 with spectinomycin, and Sanger sequenced (Azenta) using primers JCP49/50. RS wildtype or indicated mutants were made electrocompetent, transformed with pINT-ts-tet and selected for with tetracycline. Strains carrying pINT-ts-tet were made electrocompetent and transformed with appropriate pCD13PSK plasmid. Of note, following 1 h of recovery at 37°C, the culture was moved to 42°C for 30 min before plating on spectinomycin. Integration of pCD13PSK at the *attB* site was confirmed by colony PCR using JCP51/53. To additionally confirm, the region of the chromosome encoding the cloned genes and pCD13PSK with amplified with JCP51/539 and Sanger sequenced (Azenta) using primers JCP49/50. For experiments, spectinomycin was not used because the integration is stable (*42*).

### Strain construction: plasmid-borne complementation

P*_rpoE_rpoE-rseA* was cloned with JCP554/582 and assembled with pBAV amplified with JCP280/552 Plasmids were propagated in *E. coli* NEB5α, Sanger sequenced (Azenta) using primers JCP339/340, and electroporated into appropriate strain with kanamycin selection.

### Quantitative PCR

For LL-37 treatment: RS was grown overnight (16-18h) in MHB, subsequently 6 µl of culture was added to fresh 3 mL MHB and grown to OD_600_ of approximately 0.3. 50 µg/mL human cathelicidin LL-37 (Elabscience, E-PP-1466) was added for 30 minutes. 100 µl aliquots were taken before and after LL-37 exposure and plated on MHA following serial dilution.

RNA was isolated using RNeasy Mini Kit (Qiagen #74104) then treated with DNAse (BioLab #M0303S) according to manufacturer’s instructions. RNA was stored at -80 °C.Reverse transcriptase quantitative PCR was performed with Power SYBR Green RNA to CT 1-Step Kit (Applied Biosystems #4389986). *cys*G served as the housekeeping gene for normalization. *cysG* (JCP241/242), *mgtA* (mgtA-1F/R)*, arnB* (arnB-3F/R)*, rseA (rseA-2F/R),* and *rpoE* (JCP608/609) were detected by primers detailed in Supp. Table 5. Fold change was determined for each biological replicate by comparison to the same replicate without LL-37 exposure. Fold change was calculated using the 2^-ΔΔCT^ method(*43*).

### LL-37 killing assay

Strains RS and the Δ*rseA* mutant were streaked to MHA from -80°C glycerol stocks. After overnight growth, a colony was inoculated into 1.5 mL MHB. After approximately 16-20 h of growth, the culture was diluted 1:1000 for RS and 1:500 for Δ*rseA* into 1 mL fresh MHB, and an aliquot was taken, serially diluted, and plated on MHA for enumeration. After 3.5 h of growth an aliquot was taken for enumeration, then 100 μg/mL of LL-37 (Elabscience, E-PP-1466) was added. At 0.5, 1, 2, and 3 h following treatment, aliquots were taken for CFU enumeration.

## SUPPLEMENTAL MATERIAL

**Supplemental Figure 1.**
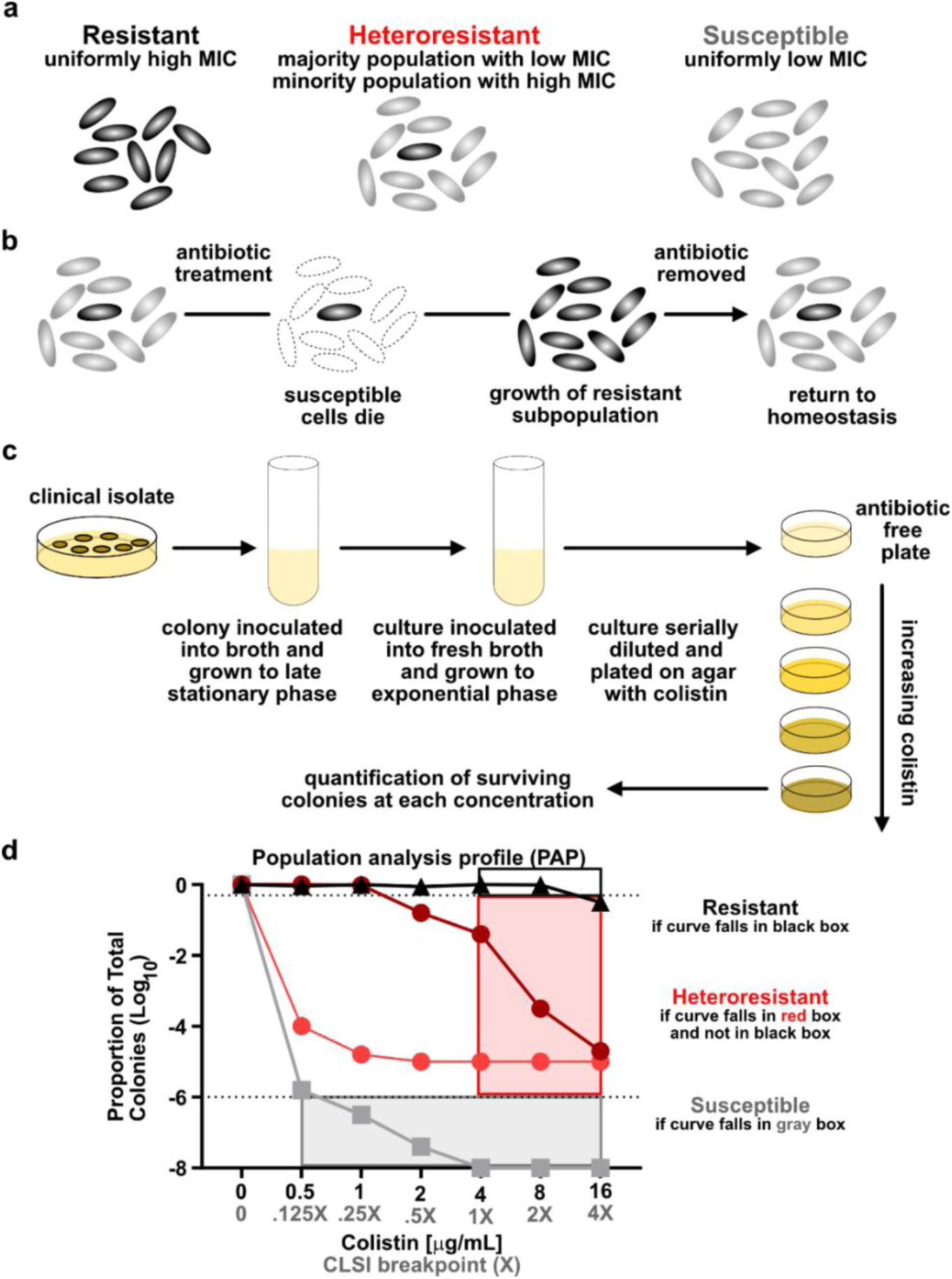
Overview of heteroresistance and population analysis profile (PAP). (a) A depiction of the cells grown from a single colony of an isolate exhibiting conventional resistance, heteroresistance, or susceptibility to a given antibiotic. (b) Population dynamics of a heteroresistant isolate following antibiotic treatment. (c) Colistin PAP: a clinical isolate is isolated and grown in broth, then subcultured into fresh broth and grown to exponential phase. The culture is then serially diluted, plated on agar with increasing concentrations of colistin, and incubated. (d) The surviving colonies are enumerated and the proportion of total colonies that survive on a given concentration is calculated: 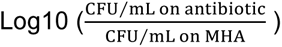. The isolate is classified as resistant if at least 50% of the total colonies grow at 1 or 2X breakpoint. An isolate is considered susceptible if less than 0.0001% (−6 logs) of the cells grow at any concentration shown. An isolate is considered heteroresistant if there is greater than 0.0001% survival at 1X and 2X breakpoint and an ≥8-fold difference between the resistance of the subpopulation and resistance of the main population. As an example, if <50% of the population survives at 1 µg/mL, the resistance of the main population is 0.5 µg/mL. The resistance of the subpopulation that survives on 16 µg/mL is >16 µg/mL, making the fold change ≥32. Along the x-axis, the breakpoints based on CLSI are shown in gray, and the corresponding concentration of colistin is in black. Because the culture density of exponential phase cultures is generally lower, in data presented in this manuscript the limit of detection is often between -5.5 to -6.5 logs. Throughout, the y-axis is set to a limit of detection for each experiment presented.

**Supplemental Figure 2.**
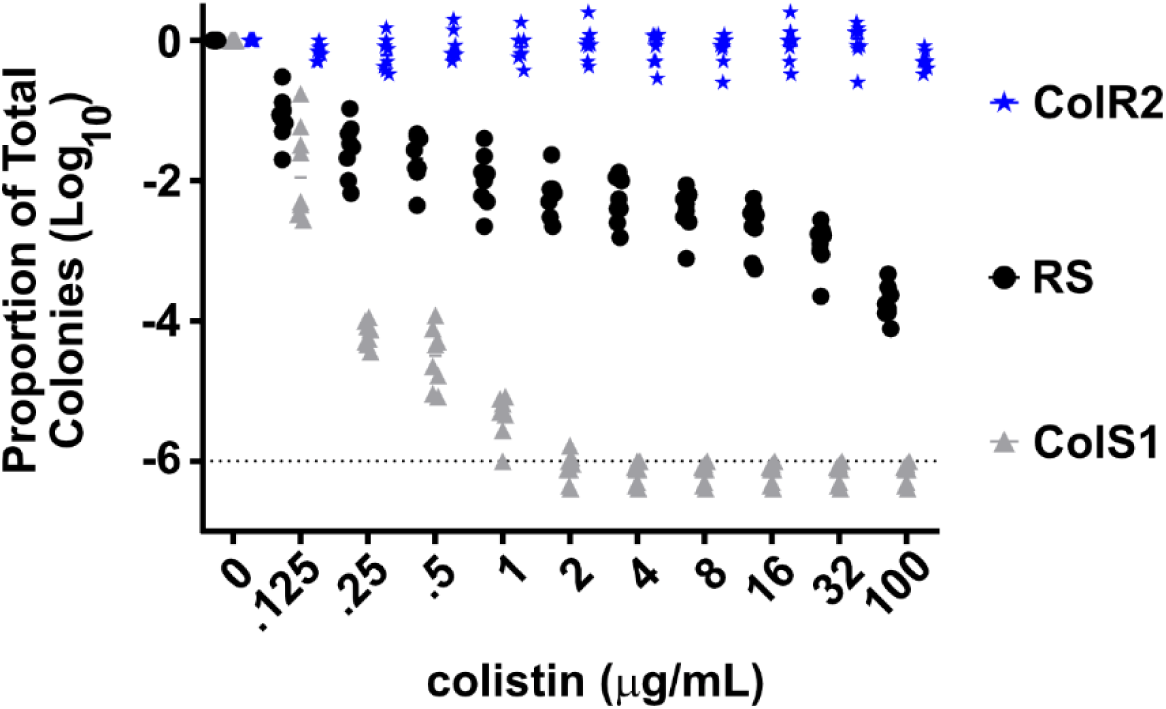
Alternative presentations of PAP of strain RS, ColS1, ColR2 from Figure 1a showing each data point, from three independent experiments with n=8 total biological replicates.

**Supplemental Figure 3.**
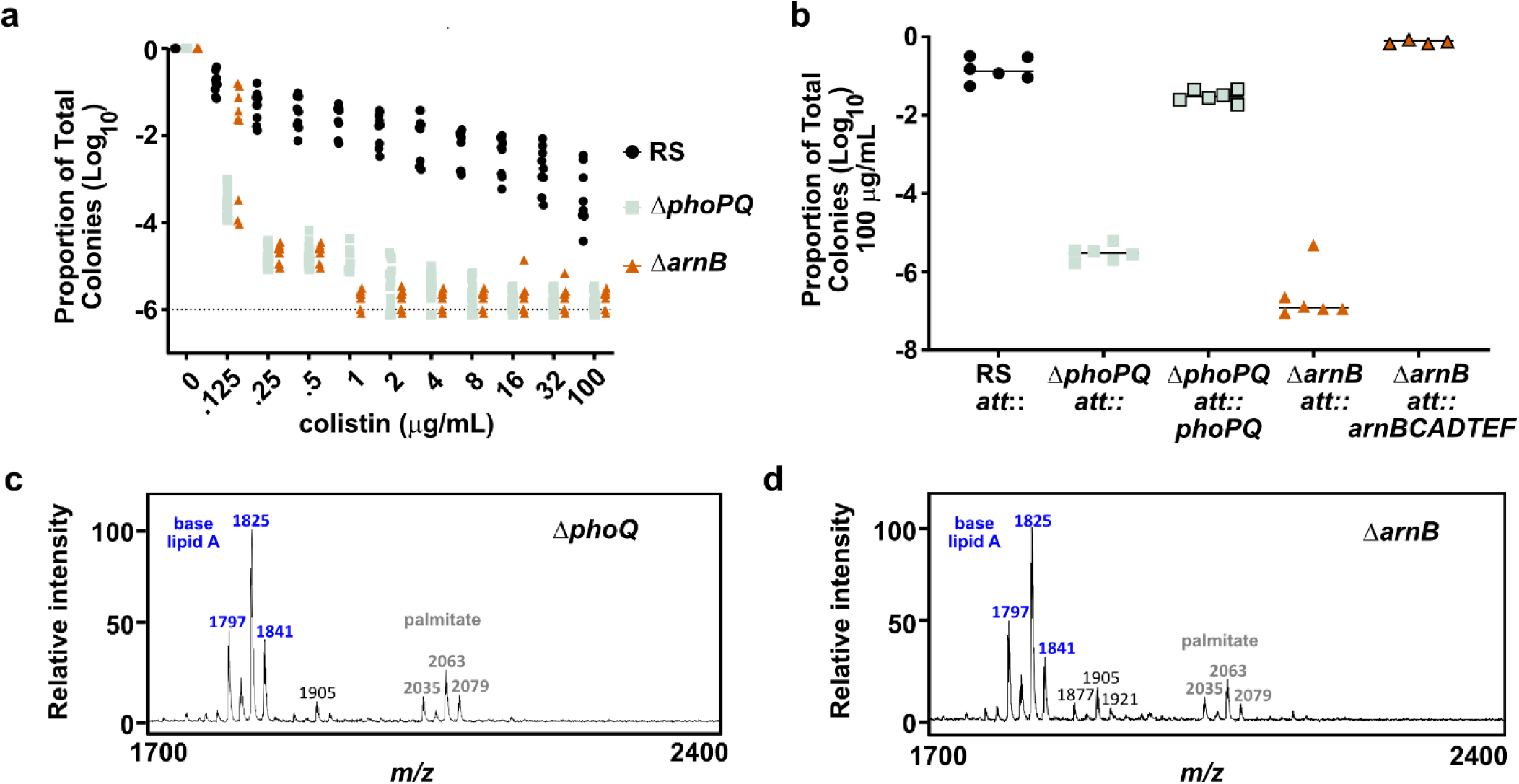
PhoPQ and ArnB are required for heteroresistance *E. cloacae* strain RS. (a) Alternative presentations of PAP of strain RS wildtype, Δ*phoPQ*, and Δ*arnB* mutants from Figure 1f on Mueller-Hinton agar (MHA) containing colistin; from three independent experiments with n=9 total biological replicates for RS and Δ*arnB*, n=12 total biological replicates for Δ*phoPQ*. (b) Proportion of the population surviving on 100 µg/mL colistin of the strains indicated, from two independent experiments n=6 total biological replicates. (c-d)) MALDI-TOF-MS spectra of strain (c) Δ*phoQ,* and (d) Δ*arnB* following growth in broth alone; a single spectra representative of 3 biological replicates is shown.

**Supplemental Figure 4.**
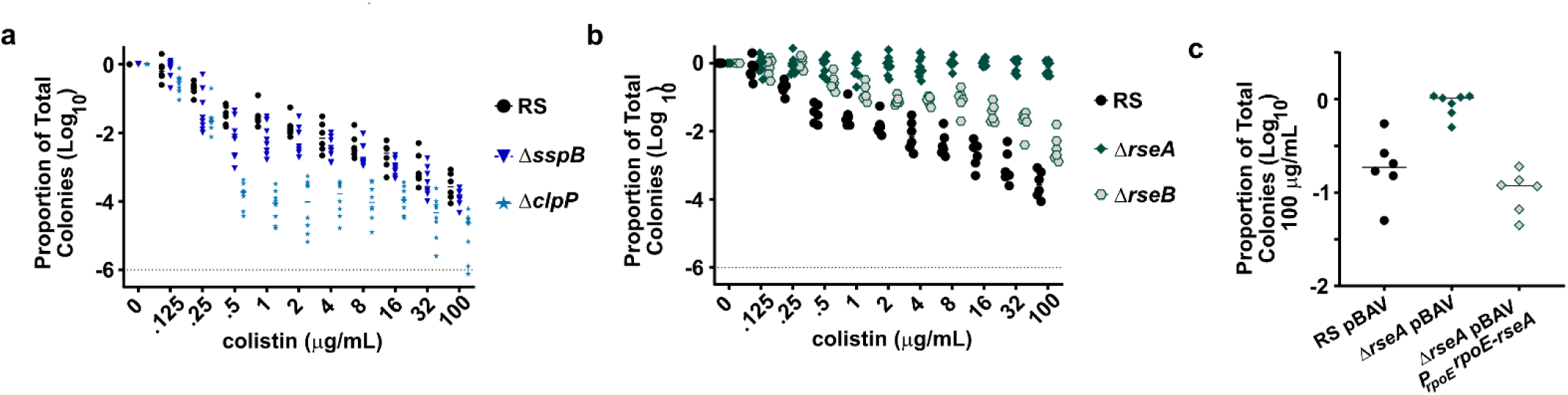
Activation of σ^E^ confers colistin resistance. (a) Alternative presentation of PAP data from strain RS and Δ*sspB* and Δ*clpP* mutants in Figure 2b. (b) Alternative presentation of PAP data from strain RS and Δ*rseA* and Δ*rseB* mutants in Figure 2c; (a-b) are from the same two independent experiments with n=6 total biological replicates for RS and Δ*rseB* and n=8 total biological replicates for the rest of the strains. (c) Proportion of the population surviving on 100 µg/mL colistin of the strains indicated, from two independent experiments with n=6 total biological replicates.

**Supplemental Figure 5.**
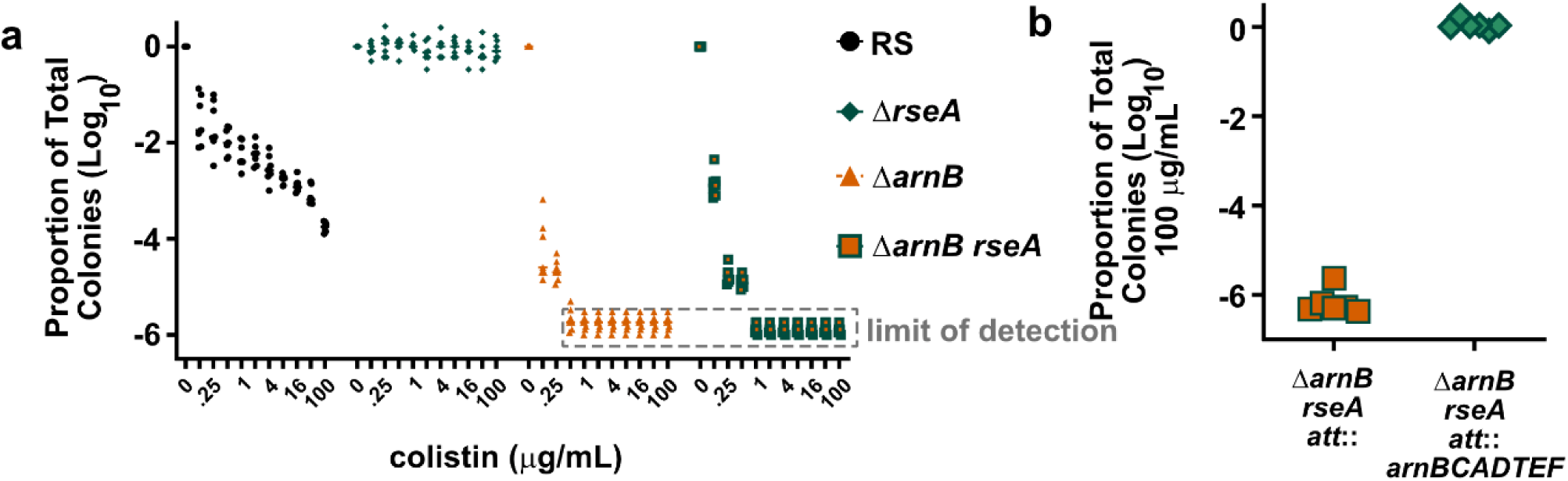
Resistance conferred by *rseA* deletion requires *arn*. (a) Alternative presentation of PAP of strain RS wildtype, and the Δ*rseA*, Δ*arnB*, and Δ*arnB rseA* mutants from Figure 4a, from two independent experiments with n=7 total biological replicates. (b) Proportion of the population surviving on 100 µg/mL colistin of the strains indicated, from two independent experiments with n=6 total biological replicates.

**Supplemental Figure 6.**
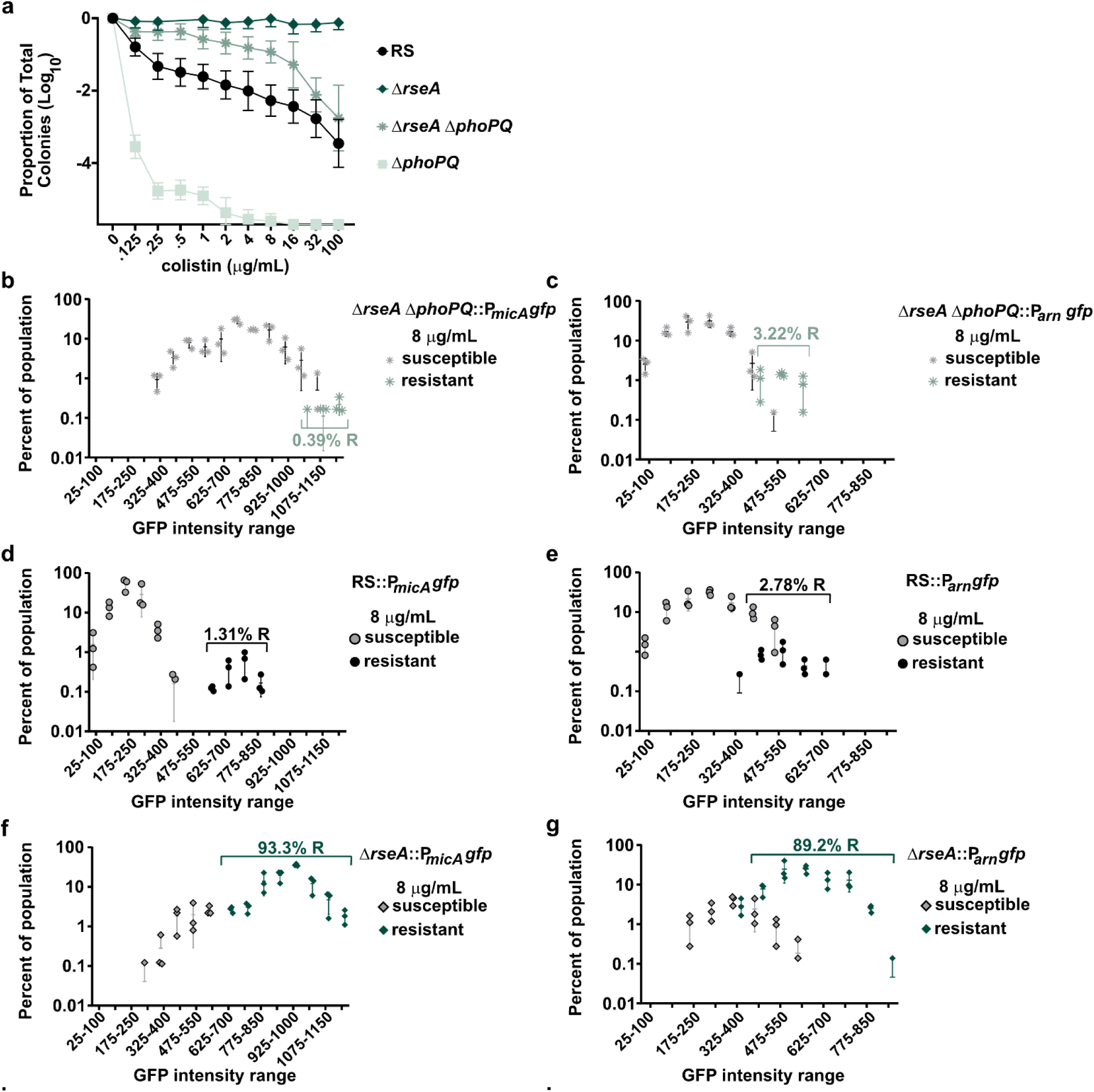
PhoPQ independent colistin resistance in Δ*rseA* background. (a) PAP of strain RS wildtype and the Δ*rseA*, Δ*phoPQ*, and Δ*rseA* Δ*phoPQ* mutants; from three independent experiments with n=9 total biological replicates for RS and Δ*arnB*, n=11 for Δ*rseA*, and n=12 total biological replicates for Δ*rseA* Δ*phoPQ* and Δ*phoPQ*. Data for RS and Δ*phoPQ* are also shown in Figure 1f. (b-g) *gfp* expression intensity, driven by the promoter indicated, in strain backgrounds as indicated, among cells prior to colistin exposure. The cell fate was monitored over time after adding colistin at the concentrations indicated, and each cell was designated as susceptible or resistant, from three independent experiments; data were determined from 700-1000 cells for each experiment. Panels d,e,f, and g are reproduced from Figures 3d, 3e, 4c and 4d, respectively, for the purpose of comparison.

**Supplemental Figure 7.**
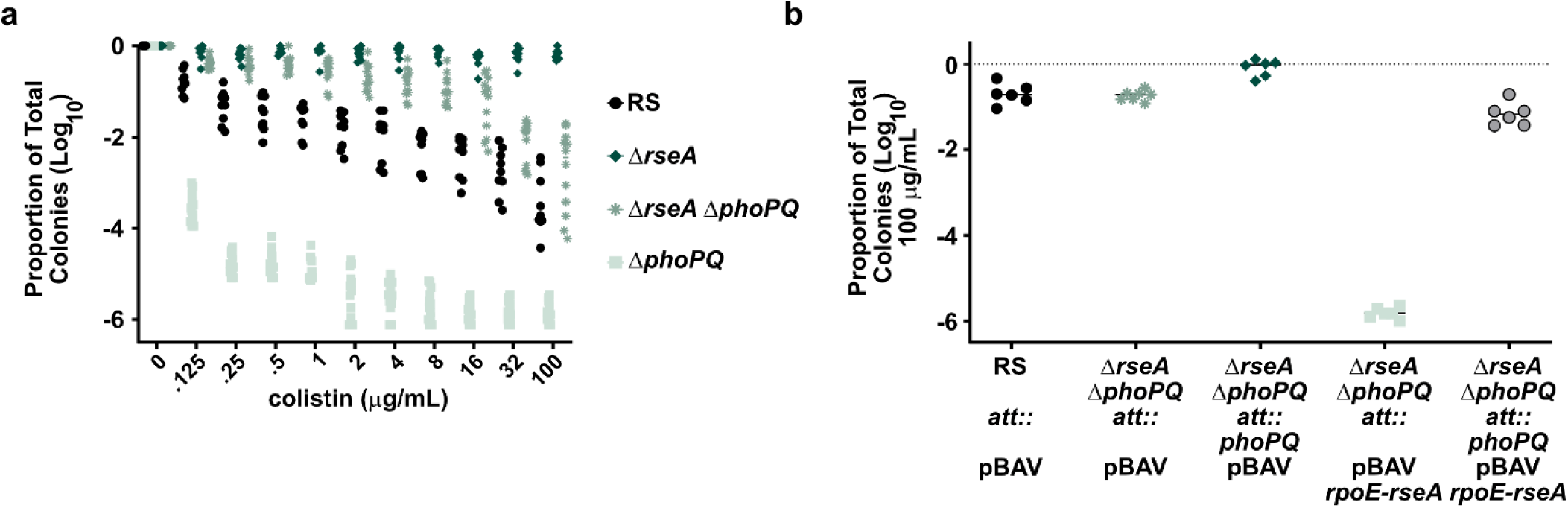
Colistin resistance in Δ*rseA* Δ*phoPQ*. (a) Alternative presentation of PAP in Supp. Fig. 8a, of strain RS wildtype and the Δ*rseA*, Δ*phoPQ*, and Δ*rseA* Δ*phoPQ* mutants; from three independent experiments with n=9 total biological replicates for RS and Δ*arnB*, n=11 for Δ*rseA*, and n=12 total biological replicates for Δ*rseA* Δ*phoPQ* and Δ*phoPQ*. Data for RS and Δ*phoPQ* are also shown in Supplemental Figure 3a. (b) Proportion of the population surviving on 100 µg/mL colistin of the strains indicated, from two independent experiments with n=6 total biological replicates, except for n=7 for Δ*rseA* Δ*phoPQ att*:: *pBAV*.

**Supplemental Figure 8.**
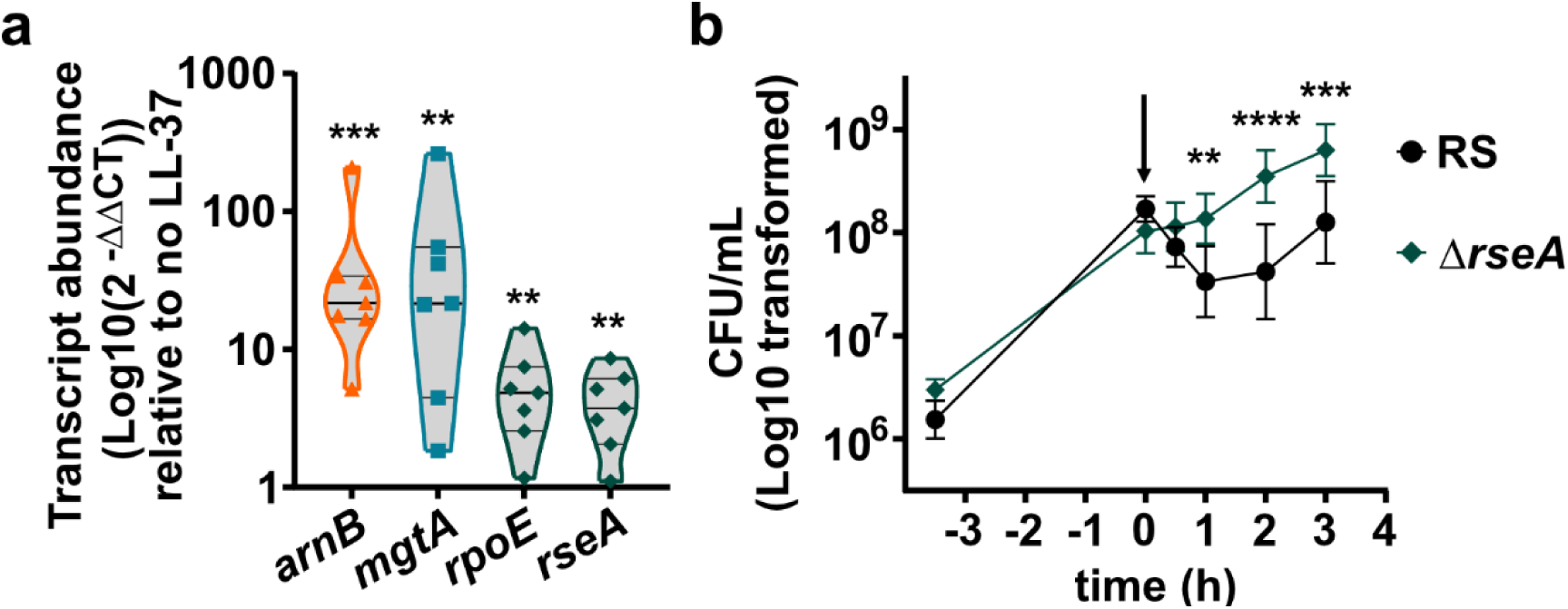
The σ^E^ pathway is activated by LL-37 and confers resistance. (a) Quantitative PCR of transcripts indicated after 30 minutes of 50 µg/mL LL-37 exposure in exponential phase cultures of strain RS wildtype, from two independent experiments with n=7 total biological replicates. *** indicates p=0.0008, ** (*mgtA*) indicates 0.0077, (*rpoE*) 0.0089, (*rseA*) 0.0084 by RM one-way ANOVA with Dunnet’s multiple comparisons correction, F (1.964, 11.78) = 18.82. (b) Culture density of strain RS wildtype and the Δ*rseA* mutant in broth. Stationary phase cultures were diluted into fresh broth and grown for 3.5 h. 100 µg/mL LL-37 was added (indicated by arrow) and culture density was measured over time. ** indicates p=0.0011, **** p<0.0001, and *** p=0.0001 by ordinary one-way ANOVA with Šídák’s multiple comparisons test, comparing strains at each time point, F (9, 60) = 12.49. From two independent experiments with n=7 total biological replicates.

**Supplemental Table 2:** theoretical m/z units of lipid A and modified lipid A.

|  |  |  |
| --- | --- | --- |
| Molecule | m/z | Calculations |
| reduction of two carbon units of the acyl chain | 1797 | 1825 – 28 (C<sub>2</sub>H<sub>4</sub>) |
| base lipid A | 1825 | 1825 base peak |
| + hydroxylation | 1841 | 1825 + 16 (OH) |
| reduction of two carbon units of the acyl chain<br>+ phosphate | 1877 | 1825 – 28 (C<sub>2</sub>H<sub>4</sub>) + 80 (phosphate) |
| + phosphate | 1905 | 1825 + 80 (phosphate) |
| reduction of two carbon units of the acyl chain<br>+phosphoethanolamine | 1920 | 1825 - 28 (C<sub>2</sub>H<sub>4</sub>) + 123 (phosphoethanolamine) |
| + hydroxylation<br>+ phosphate | 1921 | 1825 + 16 (OH) + 80 (phosphate) |
| reduction of two carbon units of the acyl chain<br>+4-amino-4-deoxy-l-arabinose | 1928 | 1825 – 28 (C<sub>2</sub>H<sub>4</sub>) + 131 (aminoarabinose) |
| + phosphoethanolamine | 1948 | 1825 + 123 (phosphoethanolamine) |
| +4-amino-4-deoxy-l-arabinose | 1956 | 1825 + 131 (aminoarabinose) |
| + hydroxylation<br>+ phosphoethanolamine | 1964 | 1825 + 16 (OH) + 123 (phosphoethanolamine) |
| + hydroxylation<br>+4-amino-4-deoxy-l-arabinose | 1972 | 1825 + 16 (OH) + 131 (aminoarabinose) |
| reduction of two carbon units of the acyl chain<br>+palmitoylation | 2035 | 1825 – 28 (C<sub>2</sub>H<sub>4</sub>) + 238 (C<sub>16</sub> palmitate) |
| +palmitoylation | 2063 | 1825 + 238 (C<sub>16</sub> palmitate) |
| + hydroxylation<br>+palmitoylation | 2079 | 1825 + 16 (OH) + 238 (C<sub>16</sub> palmitate) |
| reduction of two carbon units of the acyl chain<br>+4-amino-4-deoxy-l-arabinose<br>+palmitoylation | 2166 | 1825 – 28 (C<sub>2</sub>H<sub>4</sub>) + 131 (aminoarabinose) + 238 (C<sub>16</sub> palmitate) |
| +4-amino-4-deoxy-l-arabinose<br>+palmitoylation | 2194 | 1825 + 131 (aminoarabinose) + 238 (C<sub>16</sub> palmitate) |
| Consistent with the figure labeling, blue indicates base lipid A and its variants, and orange indicates modifications which include +4-amino-4-deoxy-l-arabinose |  |  |

**Supplemental Table 3.** Transposon screen mutant identification.

| Supplemental Table 3. Transposon screen mutant identification |  |  |  |
| --- | --- | --- | --- |
| Locus tag | name | Annotation | # of colonies identified |
| NF29_00305 |  | ferrichrome transporter | 1 |
| NF29_00575 | *murG* | UDP-diphospho-muramoylpentapeptide beta-N-acetylglucosaminyltransferase; disrupted; | 2 |
| NF29_02370 |  | hypothetical protein | 1 |
| NF29_04585 |  | 23S ribosomal RNA | 1 |
| NF29_07140 | *arnD* | 4-deoxy-4-formamido-L-arabinose-phospho-UDP deformylase | 1 |
| NF29_08180 | *sspA* | stringent starvation protein A, upstream in operon with or sspB | 1 |
| NF29_09225 |  | DNA topoisomerase IV subunit A | 1 |
| NF29_15160 |  | receptor for colicin s4 | 1 |
| NF29_16590 |  | acetylserine transporter | 2 |
| NF29_18350 |  | PEP synthetase regulatory protein | 24 |
| NF29_19185 |  | dihydroorotase | 1 |
| NF29_20475 |  | multidrug transporter | 3 |
| NF29_22440 | *clpP* | clpP, an ATP-dependent Clp protease proteolytic subunit | 1 |
| NF29_22595 |  | prepreprotein translocase subunit YajC, in operon upstream of secD | 1 |
| NF29_22635 |  | proY proline specific permease | 1 |
| **Mutants Screened** |  | **Total hits** | **Rate** |
| 16,524 |  | 42 | 0.25400% |

**Supplemental Table 4:** Strains and plasmids.

| Supplemental Table 4: Strains and plasmids |  |  |
| --- | --- | --- |
| Strains |  |  |
| Name | Description | Source |
| *Enterobacter cloacae* RS | "wildtype" | Band et al (1) |
| *E. cloacae* ColS1 | Colistin susceptible clinical isolate | Weiss lab collection (2) |
| *E. cloacae* ColR2 | Colistin resistant clinical isolate | Weiss lab collection |
| ΔphoPQ | Deletion of *phoPQ* (NF29_18845-NF29_18850) | This study |
| ΔarnB | Deletion of *arnB* (NF29_07125) | This study |
| RS::P<sub>arn</sub>*gfp* | pCD13PSK P<sub>arn</sub>*gfp*+ integrated at *attB* site | This study |
| ΔphoPQ::P<sub>arn</sub>*gfp* | pCD13PSK P<sub>arn</sub>*gfp*+ integrated at *attB* site of ΔphoPQ | This study |
| ΔsspB | Deletion of *sspB* (NF29_08180) | This study |
| ΔclpP | Deletion of *clpP* (NF29_22440) | This study |
| ΔrseA | Deletion of *rseA* (NF29_11120) | This study |
| ΔrseB | Deletion of *rseB* (NF29_11125) | This study |
| RS::P<sub>micA</sub>*gfp* | pCD13PSK P<sub>micA</sub>*gfp*+ integrated at *attB* site | This study |
| ΔrseA::P<sub>micA</sub>*gfp* | pCD13PSK P<sub>micA</sub>*gfp*+ integrated at *attB* site in ΔrseA | This study |
| ΔarnB *rseA* | *rseA*::kan_FRT and ΔarnB | This study |
| ΔrseA::P<sub>arn</sub>*gfp* | pCD13PSK P<sub>arn</sub>*gfp*+ integrated at *attB* site of ΔrseA | This study |
| RS *att*:: | pCD13PSK integrated at *attB* site of RS wildtype | This study |
| ΔphoPQ *att*:: | pCD13PSK integrated at *attB* site of ΔphoPQ | This study |
| ΔphoPQ *att*::*phoPQ* | pCD13PSK P<sub>phoP</sub>*phoPQ* integrated at *attB* site of ΔphoPQ | This study |
| ΔarnB *att*:: | pCD13PSK integrated at *attB* site of ΔarnB | This study |
| ΔarnB *att*::arnBCADTEF | pCD13PSK ParnarnBCADTEF integrated at *attB* site of ΔarnB | This study |
| ΔarnB *rseA att*:: | pCD13PSK integrated at *attB* site of ΔarnB *rseA* | This study |
| ΔarnB *rseA att*::arnBCADTEF | pCD13PSK P<sub>arn</sub>arnBCADTEF integrated at *attB* site of ΔarnB *rseA* | This study |
| ΔrseA ΔphoPQ | Deletions of *rseA* and *phoPQ* | This study |
| ΔrseA ΔphoPQ::P<sub>micA</sub>*gfp* | pCD13PSK P<sub>micA</sub>*gfp*+ integrated at *attB* site of ΔrseA ΔphoPQ | This study |
| ΔphoPQ::P<sub>micA</sub>*gfp* | pCD13PSK P<sub>micA</sub>*gfp*+ integrated at *attB* site of ΔphoPQ | This study |
| ΔrseA ΔphoPQ::P<sub>arn</sub>*gfp* | pCD13PSK P<sub>arn</sub>*gfp*+ integrated at *attB* site of ΔrseA ΔphoPQ | This study |
| ΔrseA ΔphoPQ *att*:: | pCD13PSK integrated at *attB* site of ΔrseA ΔphoPQ | This study |
| ΔrseA ΔphoPQ *att*::P<sub>phoP</sub>*phoPQ* | pCD13PSK P<sub>phoP</sub>*phoPQ* integrated at *attB* site of ΔrseA ΔphoPQ | This study |
| Plasmids |  |  |
| Name | Description | Source |
| pBAV | pBAV1K-T5-*gfp* with *gfp* excised by Hifi assembly | Choby et al (3) |
| pBAV P<sub>rpoE</sub>*rpoE-rseA* | pBAV carrying P<sub>rpoE</sub>*rpoE-rseA* without T5 promoter | This study |
| pKD46-tet | Encodes lambda red recombinase, tetracycline resistant derivative of pKD46, temperature sensitive origin | Datsenko and Wanner (4) |
| pCP20 | Encodes Flp recombinase, temperature sensitive origin | Cherepanov and Wackernagel (5) |
| pCD13PSK | Pi-dependent origin plasmid with homology to *attB* site | Haldimann and Wanner (6) |
| pINT-ts-tet | Encodes helper machinery for pCD13PSK, temperature sensitive origin | Haldimann and Wanner (6) |

**Supplemental Table 5:**
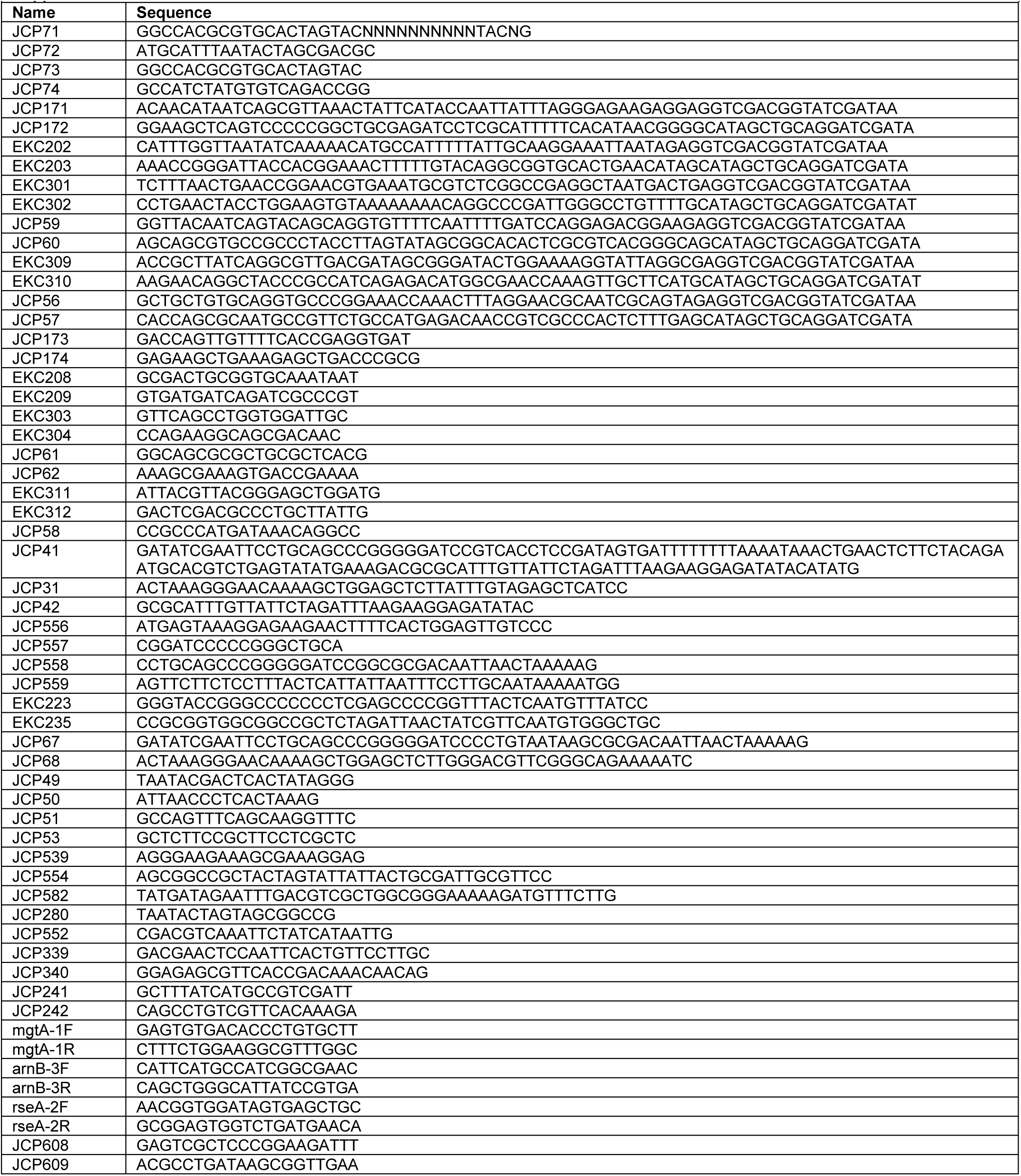
Primers.

